# OUTRIDER: A statistical method for detecting aberrantly expressed genes in RNA sequencing data

**DOI:** 10.1101/322149

**Authors:** Felix Brechtmann, Agnė Matusevičiūtė, Christian Mertes, Vicente A Yépez, Žiga Avsec, Maximilian Herzog, Daniel M Bader, Holger Prokisch, Julien Gagneur

## Abstract

RNA sequencing (RNA-seq) is gaining popularity as a complementary assay to genome sequencing for precisely identifying the molecular causes of rare disorders. A powerful approach is to identify aberrant gene expression levels as potential pathogenic events. However, existing methods for detecting aberrant read counts in RNA-seq data either lack assessments of statistical significance, so that establishing cutoffs is arbitrary, or rely on subjective manual corrections for confounders. Here, we describe OUTRIDER (OUTlier in RNA-seq fInDER), an algorithm developed to address these issues. The algorithm uses an autoencoder to model read count expectations according to the co-variation among genes resulting from technical, environmental, or common genetic variations. Given these expectations, the RNA-seq read counts are assumed to follow a negative binomial distribution with a gene-specific dispersion. Outliers are then identified as read counts that significantly deviate from this distribution. The model is automatically fitted to achieve the best correction of artificially corrupted data. Precision–recall analyses using simulated outlier read counts demonstrated the importance of combining correction for co-variation and significance-based thresholds. OUTRIDER is open source and includes functions for filtering out genes not expressed in a data set, for identifying outlier samples with too many aberrantly expressed genes, and for the P-value-based detection of aberrant gene expression, with false discovery rate adjustment. Overall, OUTRIDER provides a computationally fast and scalable end-to-end solution for identifying aberrantly expressed genes, suitable for use by rare disease diagnostic platforms.

## Introduction

No clear pathogenic variant can be pinpointed for many patients suspected to suffer a Mendelian disorder after undergoing whole exome or whole genome sequencing^1,2^. A possible reason is that the pathogenic variant is regulatory. Accurately identifying pathogenic regulatory variants is difficult. One difficulty is that, an individual harbors a very large number of rare non-coding variants, with about 60,000 non-coding single nucleotide variants compared with 475 protein-affecting rare variants per genome (with MAF < 0.005)^3^. A second difficulty is that our understanding of regulatory sequences is much poorer than our understanding of coding sequences.

Two recent studies have shown that using RNA sequencing (RNA-seq) to directly investigate gene expression defects in patients’ cells provides a promising complementary method for pinpointing pathogenic regulatory defects^4,5^. RNA-seq can help to reveal splicing defects, the mono-allelic expression of heterozygous loss-of-function variants, and expression outliers (i.e. genes aberrantly expressed outside their physiological range)^4,5^. The two studies used different approaches to identify expression outliers. Cummings *et al*.^4^ computed Z-scores on the logarithm of gene-length normalized read counts by subtracting the mean count and dividing by the standard deviation. Expression outliers were identified as read counts with an absolute Z-score greater than 3. No test of statistical significance was performed. This expression outlier analysis did not yield any convincing candidates. In contrast, the study by Kremer *et al*.^5^ identified 4 out of 6 newly diagnosed individuals as expression outliers. Read count outliers were identified as those with an absolute Z-score greater than 3 and further being statistical significant according to DESeq2, a statistical test originally developed for differential expression analyses^6^, which was applied by testing each sample in turn against the rest of the cohort. DESeq2 is based on the negative binomial distribution, which can be used to model overdispersed count data^7^. The application of a statistical test, appropriate for count data, may be one reason for the difference in success rates between the studies. However, the reason for the difference remains unclear because of the relatively small number of diagnosed cases, the absence of ground truth, and the lack of a direct comparison between the two approaches based on the same data.

The two studies did not only differ in whether or not a statistical test was applied but also differed in the way the data was corrected for confounders. Cummings *et al*.^4^ used RPKM (reads per kilobase per million mapped reads) expression values. These control for variations in sequencing depth but not for other sources of co-variation among the read counts. Controlling for further sources of co-variation is important because the identification of a gene as aberrantly expressed depends on the context, for example the sex of the donor. Genes encoded on the Y chromosome are not present, and thus not expressed, in women. However, in men, loss of the expression of a Y chromosome-encoded gene can be an aberrant expression event. Hence, not taking the sex of the donor into account would not allow for the detection of aberrantly silenced Y chromosome-encoded genes in males. While the sex of the donor is usually available and can be easily controlled for, other contexts for gene expression, such as the exact tissue of origin of the sample, the sample’s cell type composition, the genetic background, and technical biases, may not be known *a priori*, yet causing similar but less intuitive variations. Kremer *et al*.^5^ corrected expression levels for sex, biopsy site as inferred from the *HOX* gene set, and common technical sources of variation, which were identified by visual inspection of a hierarchical clustering of the samples. In a study that identified expression outliers, although not for the diagnosis of rare diseases, Li *et al*.^8^ corrected for sex and the top three genotype principal components, as well as for hidden confounding effects estimated using the probabilistic estimation of expression residuals (PEER) method^9^. However, the algorithms controlling for covariations in RNA-seq read count data used in the studies of Li *et al*.^8^ and Kremer *et al*.^5^ were neither assessed nor tuned to detect aberrantly expressed genes.

Here, we introduce OUTRIDER (OUTlier in RNA-seq fInDER), an algorithm that provides a statistical test for outlier detection in RNA-seq samples while controlling for co-variations among the gene read counts. The modeling of co-variation is performed by an autoencoder that controls for read count variations caused by factors not known *a priori*. Its parameters are optimized automatically for correcting read counts corrupted *in silico*. Autoencoders were introduced to find low-dimensional representations of high-dimensional data through a hidden layer^10–12^. They have been shown to be useful for extracting meaningful biological features from RNA-seq data^13^ and imputing missing values in single-cell RNA-seq data^14^. A subclass of autoencoders, the so-called denoising autoencoders, are used to reconstruct corrupted high-dimensional data through exploiting correlations in the data^15^. In OUTRIDER, the autoencoder approach is used to correct for the common co-variation patterns among genes. In this article we describe the OUTRIDER algorithm, its implementation, as well as its performance and results on two data sets.

## Material and Methods

### Data sets

The read counts for the rare disease cohort were downloaded from Supplementary Data 1 published as part of the study by Kremer *et al*.^5^ (https://www.nature.com/articles/ncomms15824). GTEx read counts were obtained from the GTEx Portal (V6p counted with RNA-SeQCv1.1.8, https://www.gtexportal.org/home/datasets)^16^. Read counts for the Kremer *et al*. data set were computed according to the UCSC annotation build hg19^17^, considering the full gene body. In contrast, GTEx is based on the Gencode v19 annotation^18^, and the read count of a gene is defined as the number of paired-end read pairs overlapping exons of that gene only. FPKM (fragments per kilobase per millions of reads) values were obtained using DESeq2^6^, where the gene length was defined as the aggregated length of all the exons. We then filtered for expressed genes, defined as genes for which at least 5% of the samples had a FPKM value greater than 1 (Figure S1A and B). We further filtered out samples with a sequencing depth less than the mean sequencing depth minus three standard deviations (Figure S1C and D).

### Statistical model

We assume that the read count *k_ij_* of gene *j* = 1, …, *p* in sample *i* = 1, …, *n* follows a negative binomial (NB) distribution, with gene-specific dispersion parameter *θ_j_*, and mean *μ_ij_* equal to the product of a control factor *c_ij_* and a gene-wise adjustment factor *a_j_*:

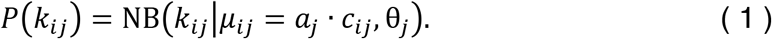

Both *c_ij_* and *a_j_* are required to be >0.01 during fitting because the NB distribution is not defined for *μ* ≤ 0. The correction factor *c_ij_* is the product of the sample-specific size factor *s_i_*, and the exponential of the sum of the co-variation factor *y_ij_* and the mean log read count per gene 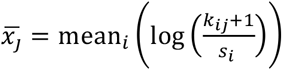:

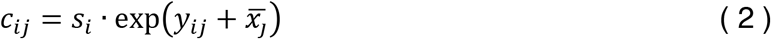

The size factors *s_i_* capture variations within the sequencing depth; they are robustly estimated as the median of the ratios of the gene read counts to their geometric means^19^. The co-variation factors *y_ij_* capture co-variations across genes; they are modeled using an autoencoder of encoding dimension *q* < *p*. Specifically,

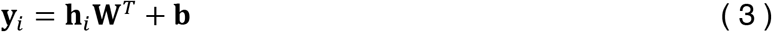

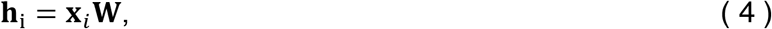

where the *p* × *q* matrix **W** is the encoding matrix and its transpose the decoding matrix, the *q*-vector **h**_i_ is the encoded representation and the *p*-vector **b** a bias term. The input to the autoencoder, **x***_i_*, is a vector of the log counts divided by the size factors centered on the gene mean:

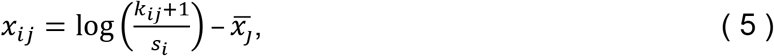

where we add one to prevent computing the logarithm of zero. In the following, we call the combination of Equations (2 - 5) the autoencoder, or short: *c_ij_* = AE(*k_ij_*).

### Fitting the autoencoder parameters

To fit the autoencoder, we set the adjustment factors *a_j_* to 1 because the goal of the autoencoder is to model as much as possible of the read count co-variation. We found that joint estimations of the dispersion parameter and the autoencoder led to poor results, possibly because one compensated for the other one. We therefore set the initial dispersion parameters for all the genes to a common value (*θ* = 25) that is arbitrary yet realistic. This value corresponds to a coefficient of variation for large read count expectations of 20%. The coefficient is also known as the biological coefficient of variation^20^. A biological coefficient of variation of 20% is typically seen for RNA-seq data sets of human cell lines with identical genetic background^21^.

For a given encoding dimension *q*, adjustment factors *a_j_*, and dispersion parameters *θ_j_*, the parameters of the autoencoder **W** and **b** are fitted by maximizing the likelihood (Equation 1) using the L-BFGS algorithm^22^ as provided by the R optimization function *optim()*. A detailed derivation of the gradient is given in the supplemental methods.

### Fitting the encoding dimension

The optimal encoding dimension is obtained by assessing the autoencoder’s performance in correcting corrupted data. We artificially introduce corrupted read counts randomly with a probability of 10^−2^ by shifting the true read counts 2 standard deviations on a log scale randomly up or down. The introduced values are rounded to the nearest integer.

With *C* denoting the set of index pairs of corrupted read counts, and 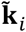 the vector of read counts for sample *i* after the injection of the corrupted read counts, an evaluation function was defined as:

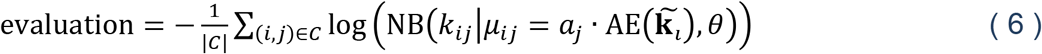

The optimal encoding dimension for this evaluation function can be selected from the integer range [1;100] by computing the loss for each value for a given *θ* = 25.

### Estimation of the dispersion and adjustment parameters

Once the autoencoder and thus the control factors *c_ij_* is fitted, the adjustment *a_j_* and dispersion *θ_j_* are obtained by maximizing the log-likelihood of Equation (1). To estimate the parameters, the negative log-likelihood is minimized for each gene independently using BFGS optimization, as implemented in the *optim()* function of R^23–26^. Initial values are estimated as 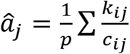 for the adjustment and 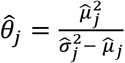 for the dispersion if 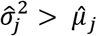, or 1 otherwise.

### P-value computation

After determining the fit parameters, the algorithm computes two-sided *P*-values using the following equation:

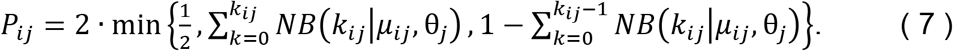

The term ½ is included to handle cases when both other terms exceed ½, which is possible because of the discrete nature of the NB distribution.

Expression levels of different genes for the same sample are correlated because of biological confounding effects such as co-regulation, which cannot be entirely excluded even after correction by the autoencoder. The computed *P*-values can therefore be correlated. Multiple testing correction was performed using the Benjamini-Yekutieli false discovery rate (FDR) method, which holds under positive dependence^27^.

### Z-score computation

Z-scores *Z_ij_* are computed on a logarithmic scale, as follows:

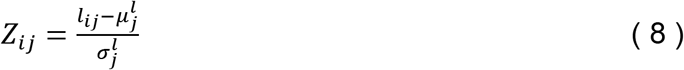

where *l_ij_* is the log_2_ corrected count calculated as 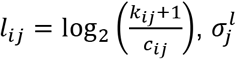 is the standard deviation of *l_ij_* for gene *j*, and 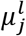 is the mean of *l_ij_* for gene *j*.

### Controlled counts

To obtain comparable counts across samples after controlling for various effects, we compute the following controlled counts: 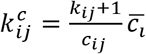. These are all of the same magnitude, independent of the control method applied. When *c_ij_* are size factors, the mean of all *c_ij_* is approximately 1 and the count is of the same magnitude as the raw count 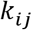. When *c_ij_* is the estimated mean, this equation provides a similar controlled count.

### Benchmark by injection of outliers

To assess the sensitivity and specificity of alternative outlier detection methods, artificial read count outliers were injected with pre-specified amplitudes (Z-scores). This process was separate from the injection of corrupted data described earlier in “Fitting the encoding dimension.” The outlier injection scheme described in this section was used to independently assess OUTRIDER’s performance in comparison with the Z-score approach.

To obtain a data set with samples that contained only a few outliers, all GTEx samples from the skin tissue not exposed to the sun were fitted and tested and only the non-outlier samples as described above in which fewer than 0.1% of all the genes were aberrantly expressed were used for the benchmark set. The read counts were then replaced with a probability of 10^-4^ by an outlier read count 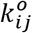, defined as:

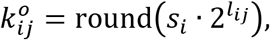

where

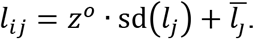

### Implementation

OUTRIDER is implemented as an R package that is available through GitHub (https://github.com/gagneurlab).

## Results

We considered two RNA-seq data sets, which we refer to as the Kremer and the GTEx data sets. The Kremer data set contains 119 RNA-seq samples from skin fibroblasts of patients with a suspected rare mitochondrial disorder^5^. This data set was analyzed in a previous study with the systematic effects corrected by manual inspection of sample correlation matrices^5^. In this study 4 genes were identified as aberrantly expressed out of 6 pathogenic genes detected by RNA-seq analysis and validated by functional assays^5^. This data set therefore served as our benchmark data set for rare disease applications. The GTEx data set contains 271 RNA-seq samples obtained from the not sun exposed skin tissue in the Genotype-Tissue Expression project (GTEx V6p^16^). We focused on these skin samples because the tissue was similar to the tissue of origin of the Kremer data set. The donors of the GTEx samples did not suffer from any condition and were not under treatment. Nevertheless, aberrant gene expression in these samples has been reported^8^.

For both data sets, we filtered out genes not expressed across the whole data set, resulting in 10,559 genes in the Kremer set and 18,250 genes in the GTEx set (Figure S1A and B). In the GTEx set, we additionally filtered out two samples because of a low sequencing depth, resulting in 269 samples (Figure S1C and D). Both data sets exhibited a strong correlation structure with very distinct sample clusters (Figure 2A and Figure S2A). These correlations may have arisen from biological variations such as the sex of the donor, the origin of the tissue, population structure, or from hidden confounders such as poorly understood systematic technical variations^5,8^. Applying the autoencoder on the read counts allowed co-variations to be estimated and corrected for. The dimension of the autoencoder was fitted using a scheme in which artificially corrupted read counts were injected and presented as the input to the autoencoder, maximizing the likelihood of the original, uncorrupted data. Based on the GTEx data set, we obtained 20 as optimal encoding dimension while 13 was optimal for the Kremer data set. We note that encoding dimensions close to the optimum yielded similar results. After the correction was applied, the co-variation clusters disappeared from both data sets (Figure 2B and Figure S2B).

**Figure 1:**
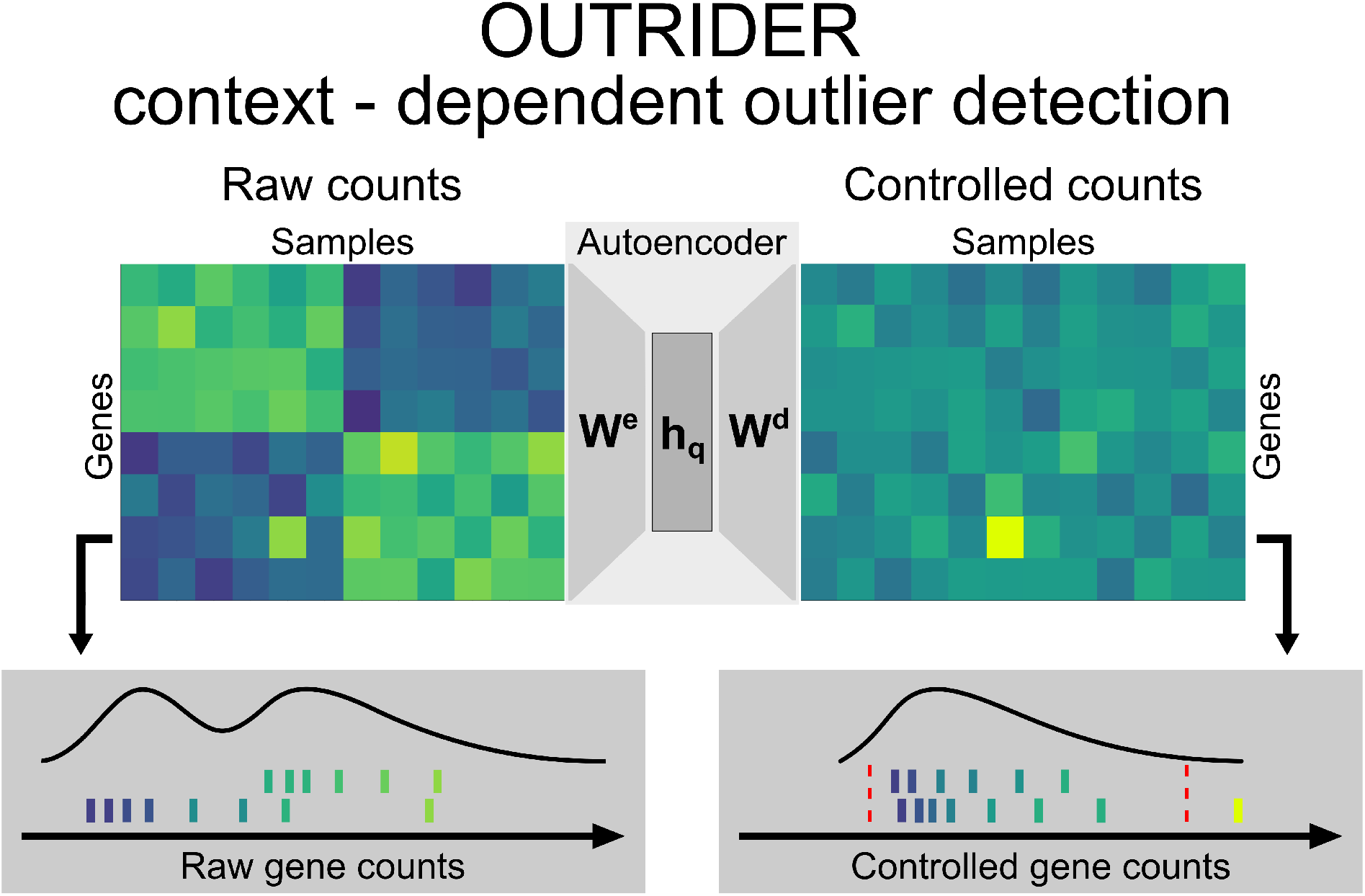
Context-dependent outlier detection. The algorithm identifies gene expression outliers whose read counts are significantly aberrant given the co-variations typically observed across genes in an RNA sequencing data set. This is illustrated by a read count (left panel, fifth column, second row from the bottom) that is exceptionally high in the context of correlated samples (left six samples) but not in absolute terms for this given gene. To capture commonly seen biological and technical contexts, an autoencoder models co-variations in an unsupervised fashion and predicts read count expectations. By comparing the earlier mentioned read count with these context-dependent expectations, it is revealed as exceptionally high (right panel). The lower panels illustrate the distribution of read counts before and after applying the correction for the relevant gene. The red dotted lines depict significance cutoffs.

**Figure 2:**
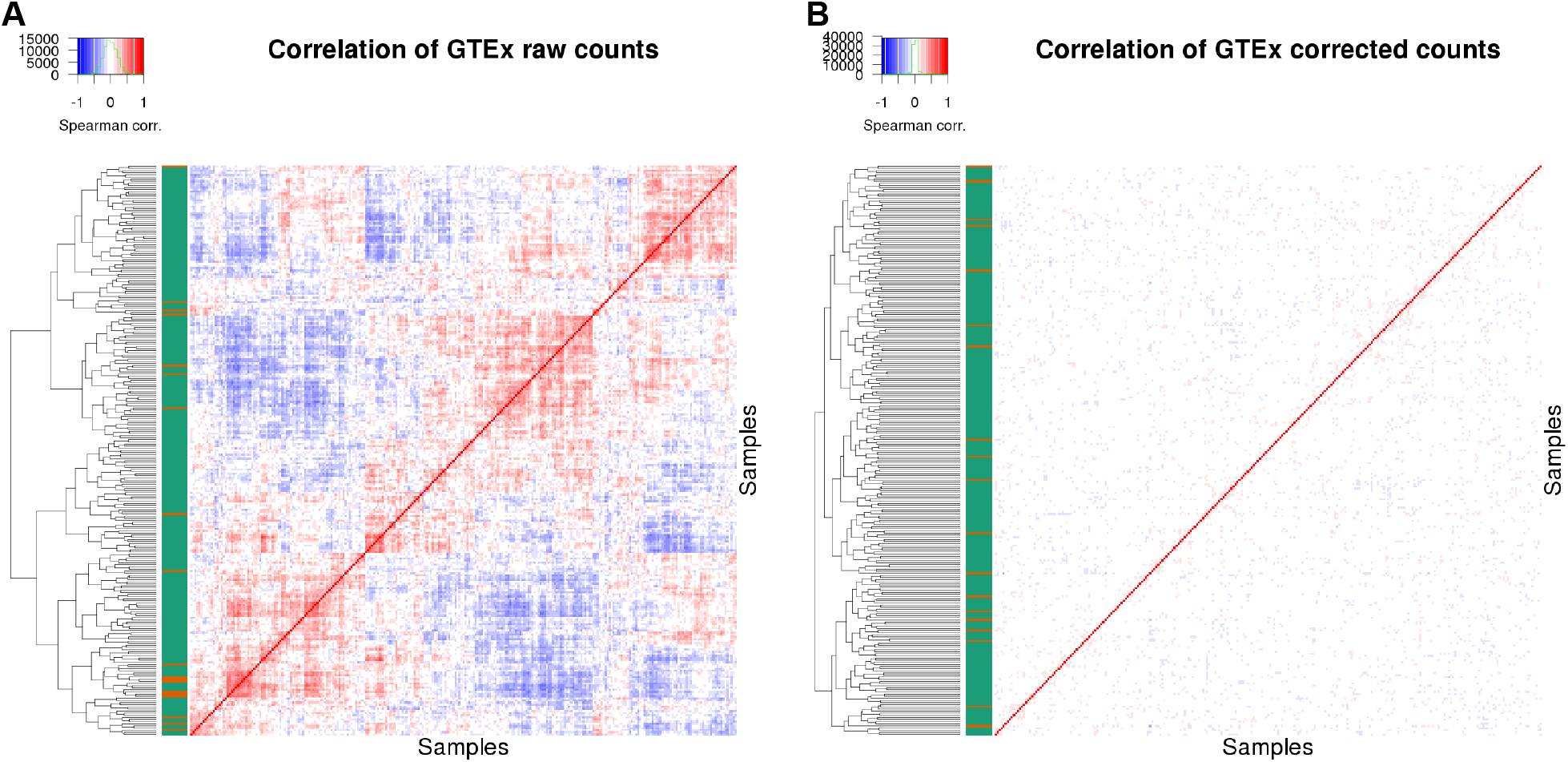
Controlling count data with autoencoders. (A) Correlation matrix of row-centered log-transformed read counts for the skin samples not exposed to the sun from the GTEx data set (of 269 samples and 18,250 genes). Red indicates a positive correlation and blue a negative correlation. The color code on the left indicates a normal sample as green and an outlier sample as orange. The dendrogram represents the sample-wise hierarchical clustering. (B) Same as for A, but with the autoencoder controlled read counts.

### Testing for aberrant expression revealed aberrant samples in both data sets

The OUTRIDER algorithm models RNA-seq read counts as a NB distribution with a mean that is the expected value provided by the autoencoder. Expression outliers are detected as significant deviations of the observed read counts from these expected values, assuming negative binomial distributions with gene-specific dispersions. Quantile–quantile plots for individual genes indicated that OUTRIDER reasonably modeled the count distribution (Figures 3B and D and S3) and that the resulting *P*-values can be used to detect outliers (Figure 3).

**Figure 3:**
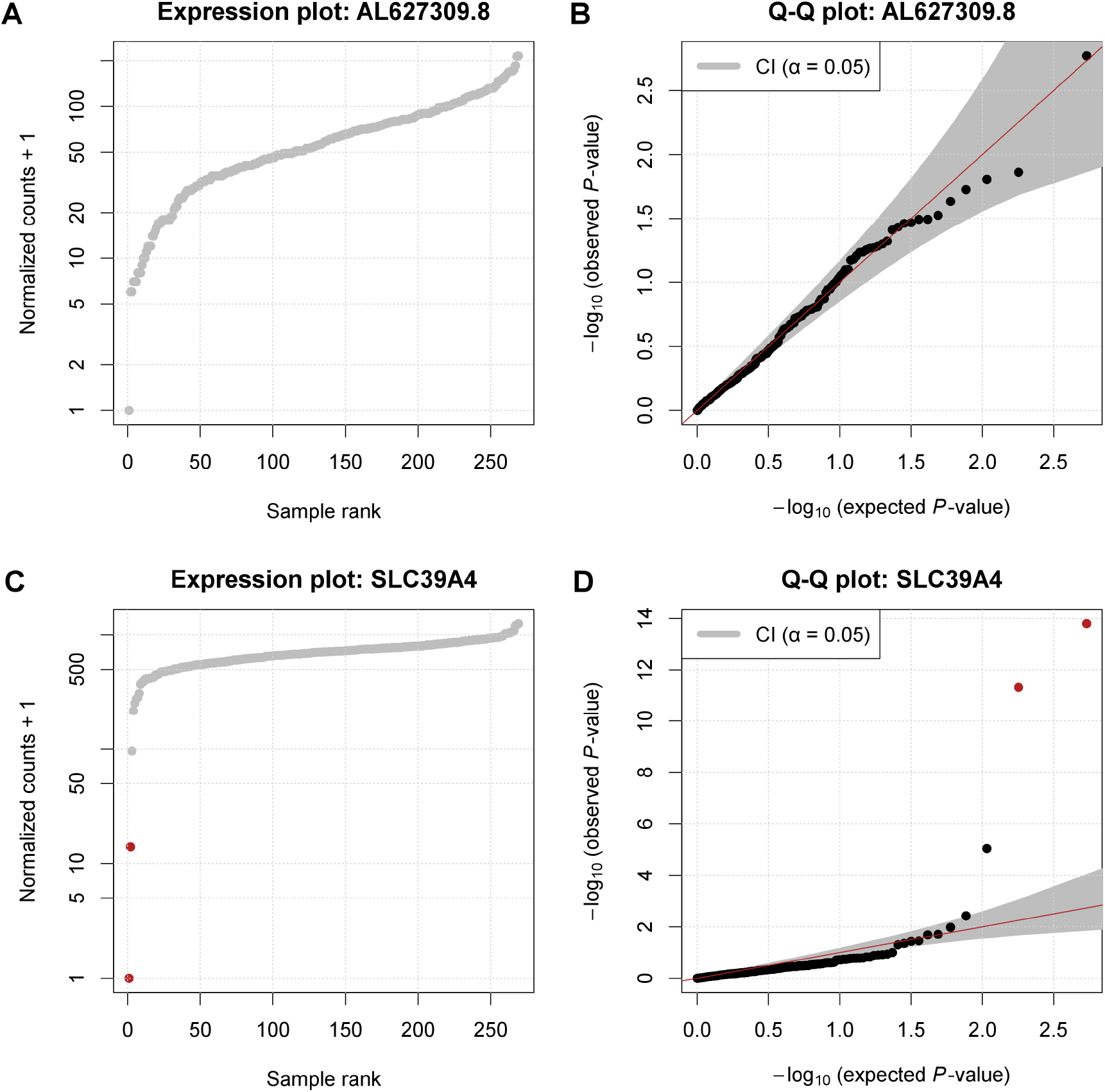
Using the negative binomial distribution for significance assessment. **(A, C)** Normalized RNA sequencing read counts plotted against their rank. **(B, D)** Quantile–quantile plots of observed *P*-values against expected *P*-values with confidence area at 5% significance level. Outliers are shown in red (FDR < 0.05). Panels A and B show data for the gene *AL627309.8* with no detected expression outlier, and panels C and D show data for the gene *SLC39A4* with two expression outliers.

Globally, at an FDR threshold of 0.05 (Benjamini–Yekutelli method^27^), we detected aberrantly expressed genes in 168 and 63 samples for the GTEx and Kremer data sets, respectively (Figure 4A and B), with 90th percentiles of 10 and 4 aberrant genes per sample. The number of significant outliers exhibited a skewed distribution across samples in both data sets, consistent with the results of Kremer *et al*.^5^. This suggested that some samples differed from others to a considerable extent. This may have been due to strong technical artifacts in the library preparation or because the samples were from donors that were genetically distant from the others or who had diseases with a severe regulatory impact. Regardless of the cause, it was difficult to narrow down the candidate pathogenic aberrantly expressed genes in these samples to a short, practical list (Figure 4C and D). We therefore introduced a cutoff (number of outliers > 0.1% of expressed genes) to filter out these samples. This resulted in 19 outlier samples (7.1%) in the GTEx data set and 4 outlier samples (3.4%) in the Kremer data set, as well as much improved global quantile–quantile plots (Figure 4E and F).

**Figure 4:**
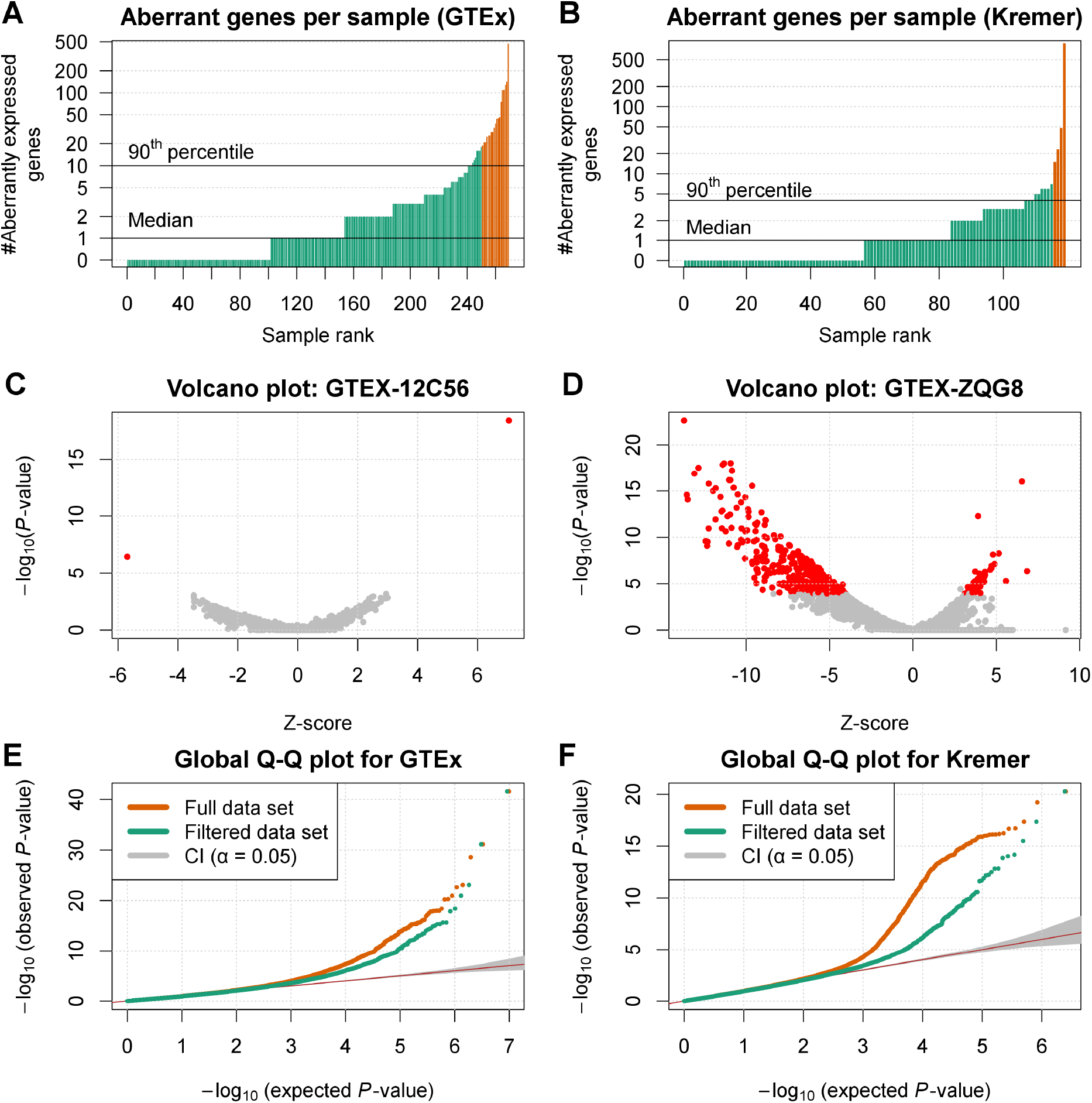
RNA sequencing expression outlier detection. **(A, B)** Number of aberrant genes (FDR < 0.05) per sample for the GTEx (A) and Kremer data sets (B). The orange bars represent abnormal samples (with >0.1% aberrantly expressed genes). **(C, D)** *P*-values versus Z-scores for a representative normal sample (C) and a representative abnormal sample (D). Genes with significantly aberrant read counts are marked in red. **(E, F)** Quantile–quantile plots for the data shown in panel A (E), and in panel B (F). Observed *P*-values are plotted against the expected *P*-values with confidence area.

### Recall benchmark

To benchmark OUTRIDER, we injected outlier read counts and tested the fraction that could be recalled. We used the 269 skin tissue samples not exposed to the sun from the GTEx data set, from which we removed 19 outlier samples. Aberrant counts were then injected into this count matrix of 250 samples and 18,250 genes. Using a frequency of 10^-4^, we injected 434 aberrant counts to create three scenarios: i) all underexpressed, ii) all overexpressed, and iii) 50%overexpressed and 50% underexpressed. Each scenario was repeated for three different simulated amplitudes (with Z-score values of 3, 4, and 6).

We then ran OUTRIDER, controlling either for sequencing depth alone or for co-variations using the autoencoder. Thereby, we monitored the recall of injected read count outliers and the precision, i.e. the number of injected outliers among the reported outliers using the different detection methods (Figure 5). Note that, the precision and recall in this setup were underestimated because the original data also contained genuine outliers. We observed that the proposed correction strategy outperformed controlling only for sequencing depth in almost all simulations. The size factor approach performed better only for very extreme overexpressed outliers (|Z| = 6). However, in this case, the recall of OUTRIDER with the autoencoder decreased by less than 5% compared to using sequencing depth alone.

**Figure 5:**
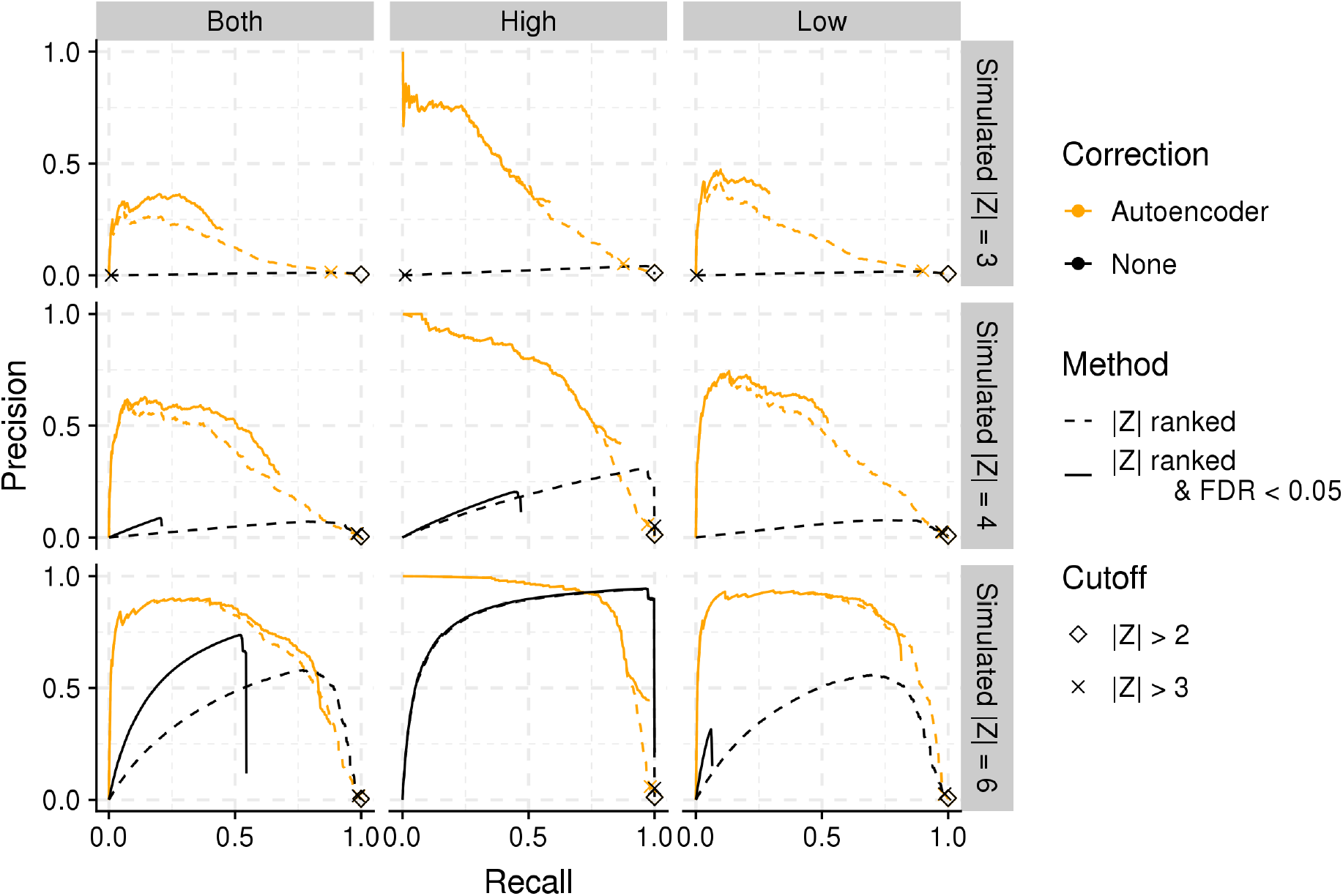
Outlier detection benchmark. The proportion of simulated outliers among reported outliers (precision) plotted against the proportion of reported simulated outliers among all simulated outliers (recall) for decreasing Z-score and for different methods. Plots are provided for three simulated amplitudes (by row, with simulated absolute Z-scores of 3, 4, and 6, top to bottom, respectively) and for all combined results (left column), for the subset of aberrantly high counts (middle column), and for the subset of aberrantly low counts (right column). The read counts were either controlled for sequencing depth only (black) or were also controlled for gene co-variation with the autoencoder (orange). The solid lines show the results restricted to significant counts only (FDR < 0.05) and the dashed lines show the results for all counts. Two commonly used Z-score cutoffs for outlier detection^4,8^ are marked with a diamond (|Z| > 2) and a cross (|Z| > 3).

The two most commonly used Z-score cutoffs^4,8^ (|2| and |3|) recalled almost all the outliers (median=99%) in each simulation, yet at the cost of a high false discovery rate (precision < 0.02). In contrast, applying a significance cutoff corrected for multiple testing reduced the rate of false discoveries and hence increased the precision, which confirms the importance of a *P*-value based strategy.

In the scenario of overexpressed genes, the precision–recall curves with and without significance overlapped. (Figure 5, middle column). This indicated that Z-scores and *P*-values for high counts gave essentially the same rankings. The advantage of *P*-values is that they provide a principled way to establish a cutoff that takes statistical significance and multiple testing into account. With low gene expression levels, the overlap of the precision–recall curves was less strong (Figure 5, right column). Only genes with a sufficiently high mean expression level (mean count >80 with recall >55% and precision >60%; Figure S4B) yielded significant *P*-values to recall the injected aberrantly low outliers. This is due to the non-negative nature of counts, where even large Z-scores do not result in significant deviations from the mean expression level (Figure S4). This limited the recall because a significant fraction of genes had a mean expression level that was too low. Overall, these results demonstrated that it is important to control for co-expression and to assess significance.

Applying OUTRIDER to the Kremer data set and filtering out the outlier samples resulted in a recall of 47 events (66%) identified by Kremer *et al*. (Figure S5) based on the 48 previously undiagnosed samples. Additionally, OUTRIDER detected 33 new expression outliers of which 19 were down-regulated. OUTRIDER was able to recall 5 out of the 6 pathogenic events (3 expression outliers, 1 mono-allelic expression, 2 splicing defects) validated by Kremer *et al.*. While Kremer at al. additionally detected the mono-allelically expressed gene *ALDH18A1* as an expression outlier it was not significant using OUTRIDER. In this case, a compound heterozygous defect was detected in *ALDH18A1*, where only the missense mutation was expressed. Hence, there was a reduction of about 40% of the overall gene expression. Nevertheless, *ALDH18A1* was the gene with the second most extreme P-value among the down-regulated genes in this sample. Furthermore, *ALDH18A1* can also be detected with a test for mono-allelic expression^5^, arguably the most appropriate aberrant expression category in this case. Two further genes, *CLPP* and *MCOLN1*, diagnosed by Kremer *et al*.^5^ by testing for splicing defects, were detected by OUTRIDER as expression outliers. Those cases exhibited reduced gene expression due to severe splicing defects.

## Discussion

We have introduced OUTRIDER, a new end-to-end solution for identifying aberrantly expressed genes within RNA-seq data, controlling for hidden confounders in an automated fashion, and providing statistical significance estimates. OUTRIDER combines an autoencoder, which allows for automatic correction of technical and biological variations among genes, and a statistical test based on a negative binomial distribution. A precision-recall analysis demonstrated the advantage of correcting for co-variations and employing significance-based thresholds. OUTRIDER has three advantages over preceding methods. First, it computes *P*-values, which can be adjusted to control the false discovery rate. Z-score-based approaches lack *P*-values, so the setting of cutoffs is arbitrary. Second, the parameters of the model are automatically fitted evaluating the ability of the model to correct for simulated corrupted read counts. Third, OUTRIDER is implemented and made available as an open source R package (from https://github.com/gagneurlab/OUTRIDER). The package provides plotting functionality and allows full analyses to be made with only a few lines of code. Furthermore, the package comes along with a comprehensive vignette guiding the user through a typical analysis.

We implemented OUTRIDER so that it is not restricted to the autoencoder, allowing the outlier test to be used with any other normalization method. In particular, autoencoders with additional layers could be employed to capture nonlinear relationships. However, the analysis of correlations post-control did not suggest the need for a more complex autoencoder. This is consistent with the study of Way and Greene, who modeled co-variations in RNA-seq samples using a single-layer autoencoder^13^. Independently of the way the correction is modeled, OUTRIDER offers functionality for finding the optimal parameter set using a modeling scheme based on corrupted count data. An advantage of this hyperparameter optimization is that no manual intervention is needed to find the optimal parameters.

Despite controlling for co-variations, some samples exhibited a much larger number of aberrant genes than others in both of the data sets investigated. This is consistent with the findings of Kremer *et al.^5^*. In outlier samples where hundreds of genes are identified as aberrant, it is difficult to draw any conclusion about the cause of such aberrant expression profiles. We therefore label those samples as outlier samples and recommend to investigate them and their global expression profile changes in a separate analysis. Some of these global changes may have a technical basis, such as the two discarded samples from the GTEx data set with aberrantly low coverage, whereas others may reflect the phenotype of the disease. Alternative methods have been developed to detect such samples; these exploit global measures such as the Mahalanobis distance to measure the dissimilarity of a sample from the population and to identify aberrantly expressed gene sets^28,29^.

We have not addressed the handling of replicate samples because we do not expect them to be performed by default in diagnostic settings. The reason for this is that outliers are events that show strong effects, so replicates are not essential for detecting these types of events. If a putative disease-causing event is detected, such as an aberrantly expressed gene, follow-up experiments involving assays complementary to RNA-seq are preferred over replicates to establish the functional link of the event to the disease^1,30^. In contrast, if an RNA-seq sample is suspected to have a technical problem, a new library can be prepared, and the new data is substituted for the former. Neither of these situations results in replicate samples. When replicates are available, users can combine the *P*-values of replicate samples using Fisher’s method for combining *P*-values^31^ by assuming independence of the read counts conditioned on the expected means predicted by the autoencoder.

In general, the autoencoder controlling scheme and the read count modeling approach benefit from additional sequencing data; the more data that can be combined, the better the estimation of the typical patterns within a population will be. This holds true when the overall data is equally distributed across population structures or sequencing protocols, because each sample is assumed to be an independent representative of the whole population. This assumption was partially violated in this study because RNA-seq data sets such as GTEx comprise >85% Caucasian individuals^16^. Such overrepresentation of a given population in the data set is disadvantageous to our method, and additional samples from underrepresented groups would be especially beneficial. More testing is needed to assess whether our strategy for controlling raw read counts can control for different data sources including data from multiple sequencing platforms or control data sets. The ability to control for different protocols enables count data to be combined from multiple sources. This would allow studies with a few samples to merge their results with sources such as the publicly available GTEx data set^16^. Currently, the best practice is to use the same cell handling and library preparation protocol which reduces the analyzable data set and therefore limits the statistical power.

The initial aim of developing OUTRIDER was to create an expression outlier detection framework for RNA-seq data in a diagnostic setting. Our primary focus was to identify aberrantly expressed genes in RNA-seq samples from patients with rare diseases. OUTRIDER will be useful for the identification of potentially disease-causing genes in patients for whom current methods, such as WES and WGS, only provide variants of unknown significance. However, our approach is not restricted to such data or experiments. In principle, OUTRIDER could model any count data derived from next-generation sequencing. Previous studies that established links between rare variants and aberrant gene expression, such as the study of Li *et al*.^8^, could benefit from OUTRIDER’s refined outlier detection approach. Our approach could also be applied to data such as DNA accessibility from ATAC-seq reads. In this case, promotor regions or enhancers would be used as features instead of gene bodies. Finally, the methodology of OUTRIDER could be adapted to detect splicing outliers or outliers in proteomics or metabolomics.

## Supplemental Data

The Supplemental Data accompanying this article includes five figures and a detailed derivation of the loss function’s gradient.

## Acknowledgments

We thank Gökcen Eraslan for fruitful and inspiring discussions about the usage of autoencoders on count data. This study was supported by the German Bundesministerium für Bildung und Forschung (BMBF) through the German Network for mitochondrial disorders (mitoNET, 01GM1113C to H.P.), the E-Rare project GENOMIT (01GM1207 to H.P.), and the Juniorverbund in der Systemmedizin ‘mitOmics’ (FKZ 01ZX1405A J.G. and V.A.Y.). A Fellowship through the Graduate School of Quantitative Biosciences Munich (QBM) supports V.A.Y. and Z.A. A.M was supported by a Fellowship through the Katholischer Akademischer Ausländer-Dienst (KAAD). C.M., F.B., V.A.Y., H.P. and J.G. are supported by EU Horizon2020 Collaborative Research Project SOUND (633974). The Genotype-Tissue Expression (GTEx) Project was supported by the Common Fund of the Office of the Director of the National Institutes of Health, and by NCI, NHGRI, NHLBI, NIDA, NIMH, and NINDS. The data used for the analyses described in this manuscript were obtained from the GTEx Portal on 06/12/17 with dbGaP accession number phs000424.v6.p1.

## Author Contributions

J.G. conceived the project and overviewed the research with the help of Z.A, H.P., and V.A.Y.F.B., A.M., C.M. analyzed the data. F.B., C.M. and A.M. developed the software. D.M.B. and M.H contributed to the software development and early stage data analysis. J.G. and Z.A. devised the statistical analysis. F.B., V.A.Y., C.M., and J.G. made the figures. F.B., C.M., A.M., V.A.Y. and J.G. wrote the manuscript. All authors performed critical revision of the manuscript.

## Web Resources

OUTRIDER: https://github.com/gagneurlab/OUTRIDER

GTEx: https://www.gtexportal.org/home

Supplementary data and additional results: https://i12g-gagneurweb.in.tum.de/public/paper/OUTRIDER

## Supplemental Data

### Supplemental Figures

**Figure S1:**
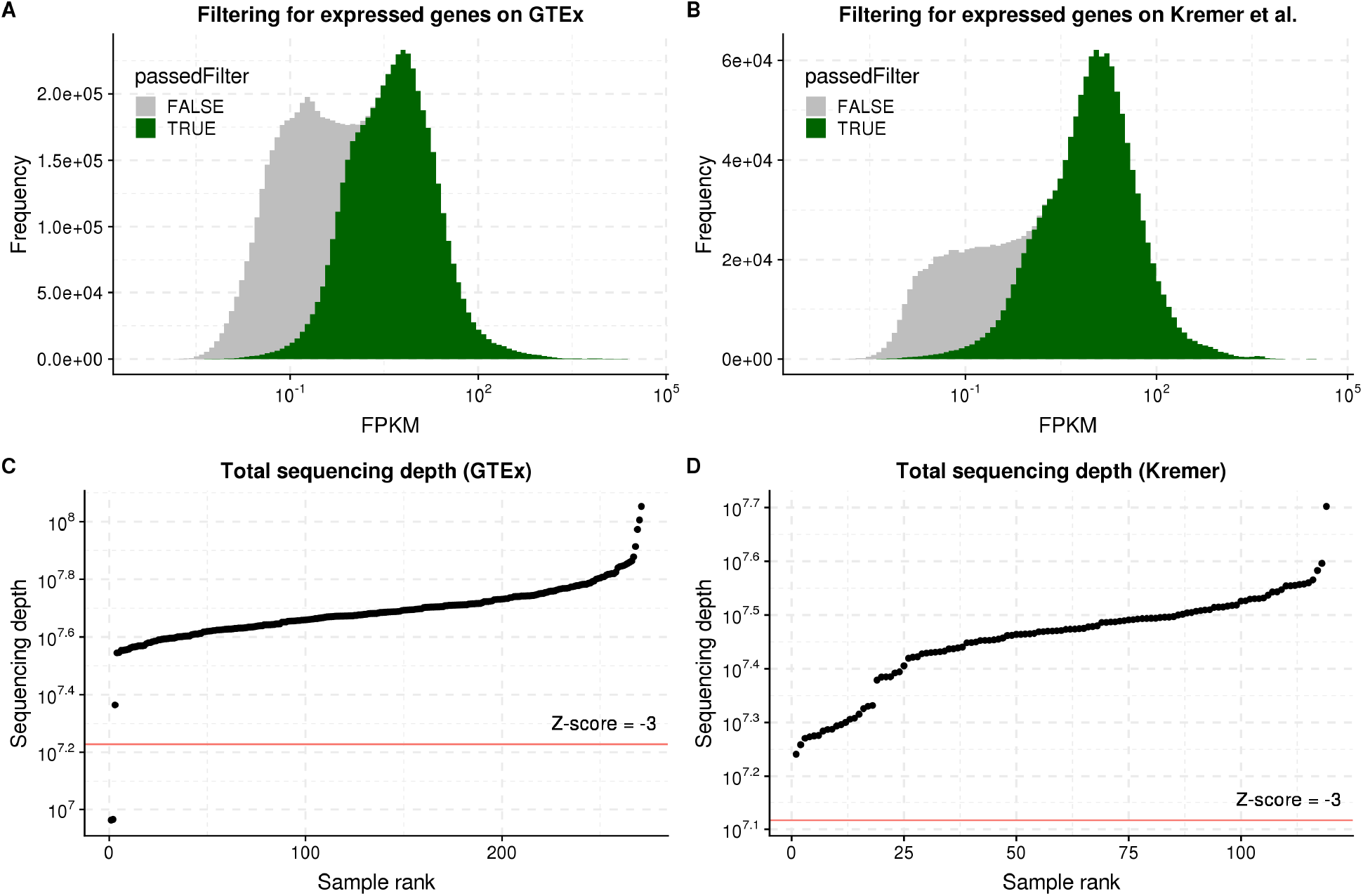
Filtering of genes and samples. **(A)** Histogram of the RPKM values for the GTEx data set grouped according to the filter status. Green indicates the genes that passed the filter and gray those that were filtered out. **(B)** The same as A, but for the Kremer data set. **(C)** The total sequencing depth sorted from low to high for all the GTEx samples. The red horizontal line represents the Z-score cutoff of -3. **(D)** The same as C, but for the Kremer data set.

**Figure S2:**
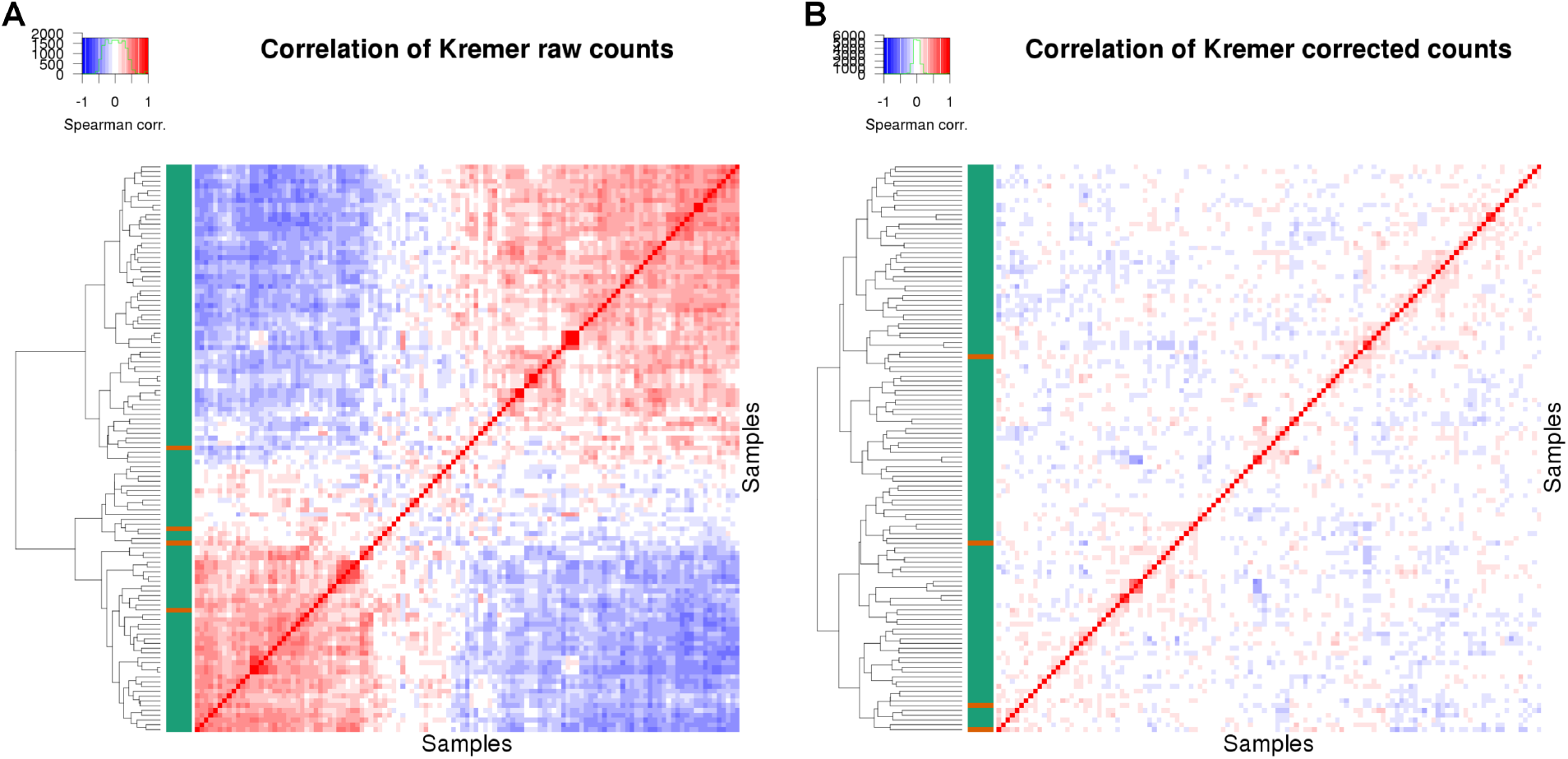
Controlling count data with autoencoders. **(A)** Correlation matrix of row-centered log-transformed read counts for the Kremer data set (with 119 samples and 10,559 genes). Red indicates a positive correlation and blue a negative correlation. The color code on the left indicates a normal sample as green and an outlier sample as orange. The dendrogram represents the sample-wise hierarchical clustering. **(B)** Same as in A, but with the autoencoder controlled read counts.

**Figure S3:**
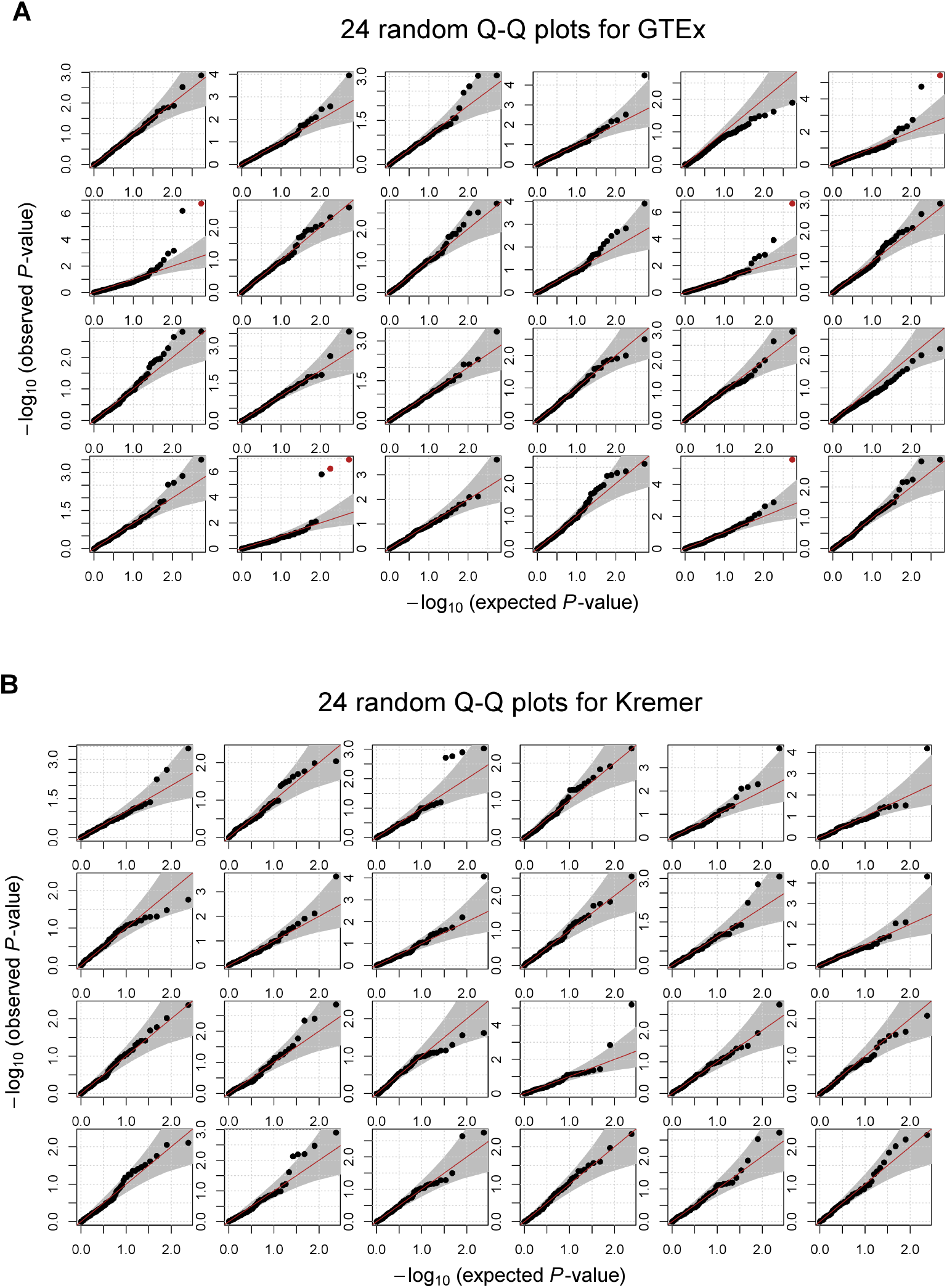
Using the negative binomial distribution for read counts. **(A)** Quantile–quantile plots for 24 randomly selected genes from the GTEx data set. **(B)** Same as in A, but for the Kremer data set.

**Figure S4:**
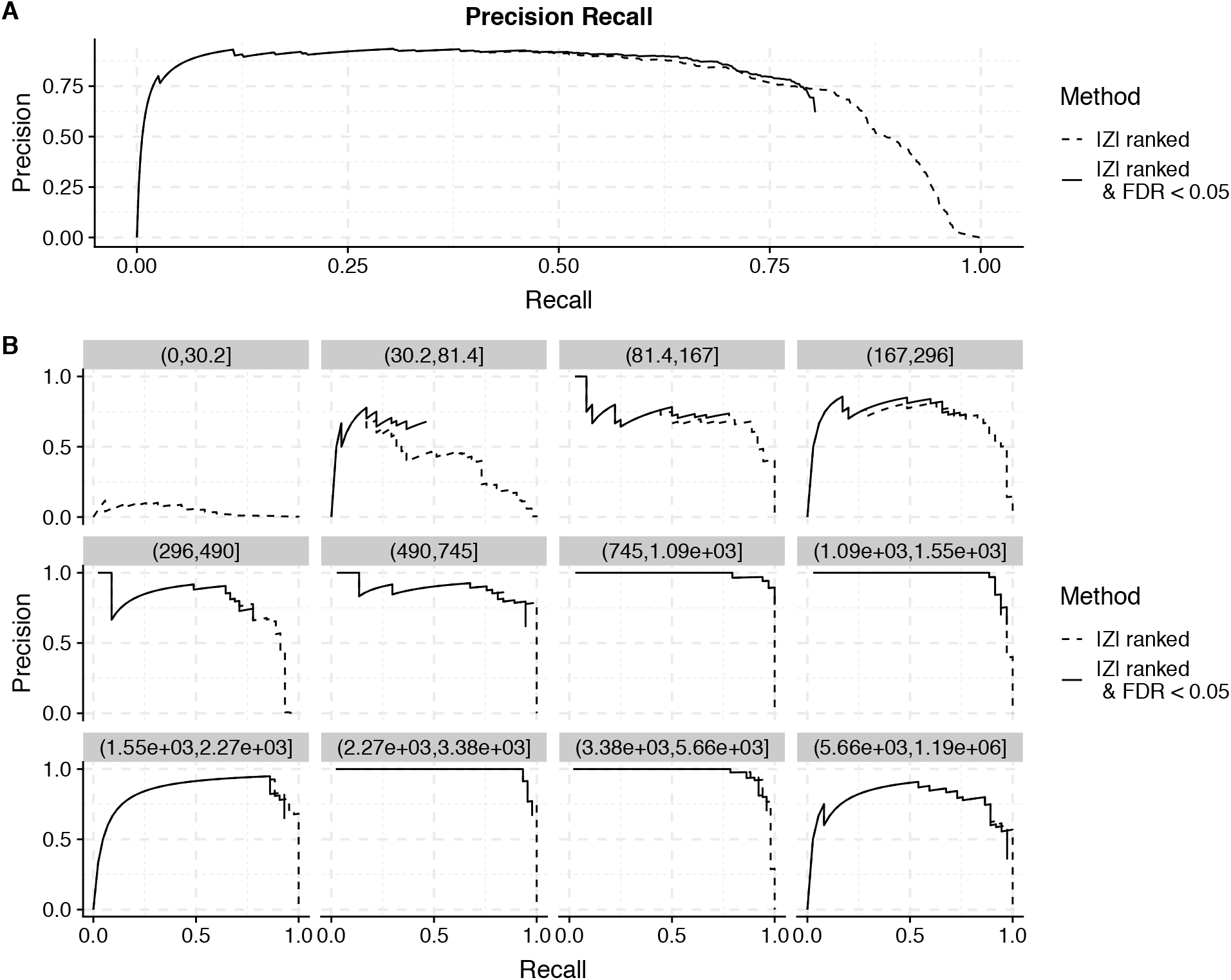
Expression level-dependent recall. Precision versus recall for artificially injected low expression outliers with a Z-score of 6. **(A)** For all the injected outliers. **(B)** Split into 12 equal bins according to the mean expression level of the genes. Only a small fraction of the injected outliers was significant for the two lowest bins, with a mean expression level smaller than 81 counts.

**Figure S5:**
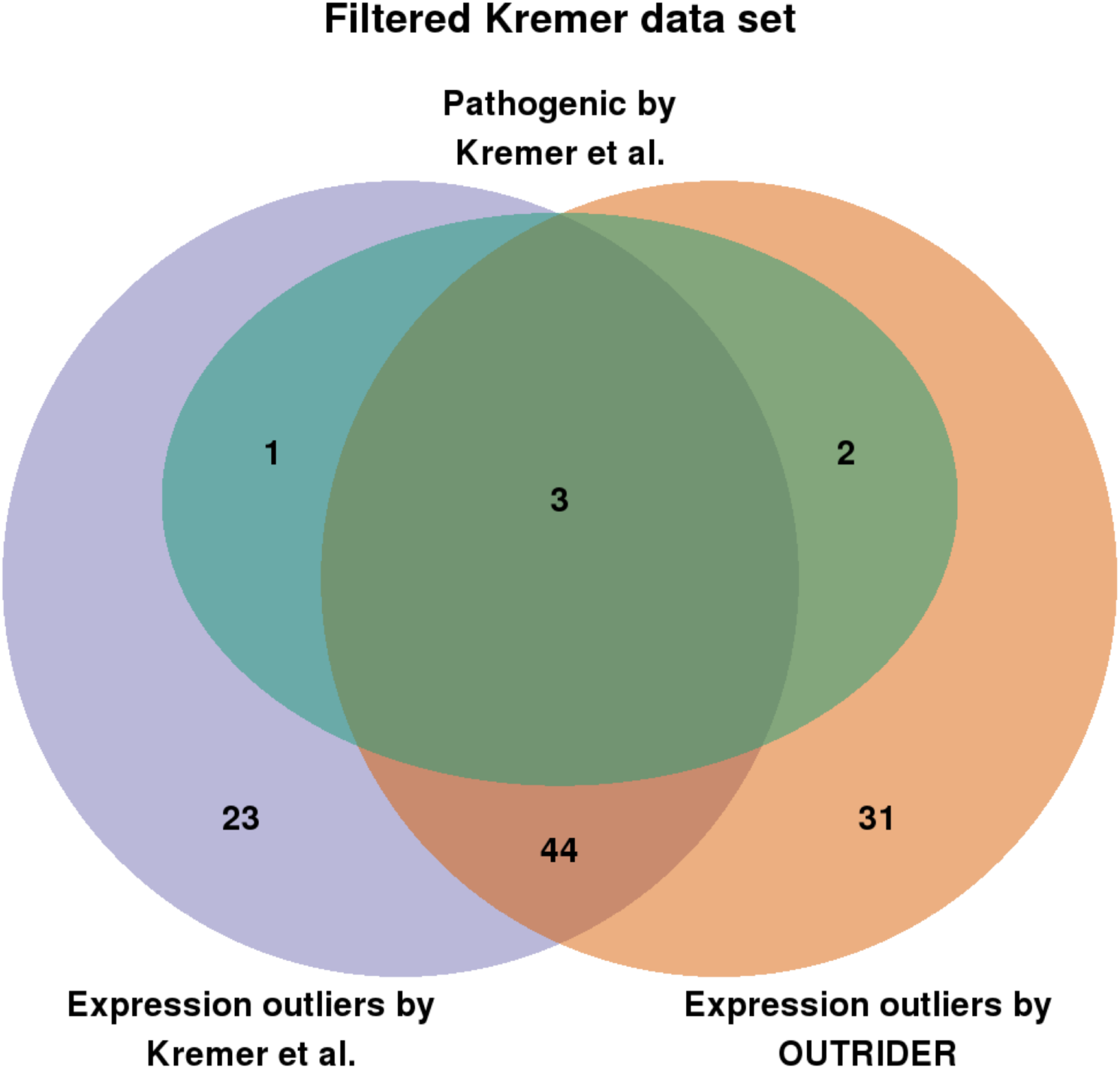
Benchmark of OUTRIDER using the Kremer data set. Venn diagram of the expression outliers detected by OUTRIDER (orange), expression outliers detected by Kremer *et al*. (violet), and pathogenic events validated by Kremer *et al*. (green) within 48 previous to Kremer *et al*. undiagnosed samples^5^.

### Supplemental Methods

We use L-BFGS to fit the autoencoder model as described in the Methods. To speed up the fitting we implemented the gradient as derived below.

The expectations *μ* arc modeled by:

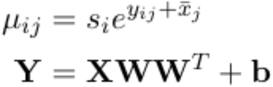

where the matrix **X** is given by the matrix: 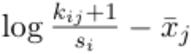. The negative binomial log likelihood is given by:

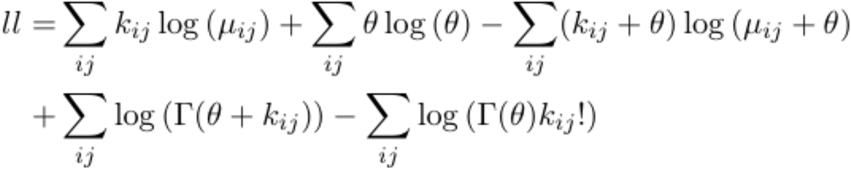

For the derivation of the gradient only the first and third term need to be considered, as all other terms are independent of *μ*.

Computing the derivative of the first term with respect to the matrix **W** by subsituting the autoencoder model for *μ*. Here the operations log [**A**] and exp [**A**] are understood to be element-wise for a matrix or vector **A**,

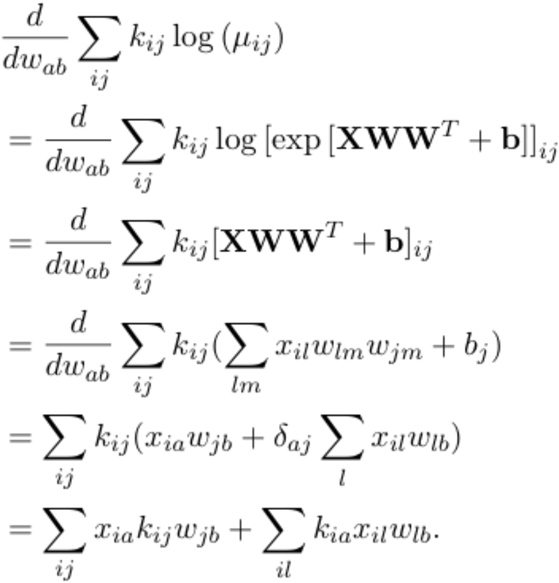

Which can be written as:

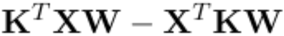

Equivalently the derivative of the third term is:

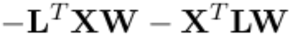

where the components of the matrix L are computed by:

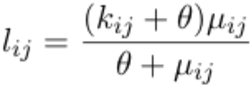

The combined result is then:

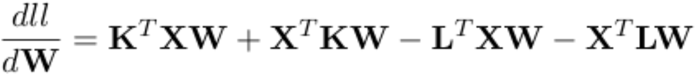

The derivative of the first term with respect to the bias **b** is computed as:

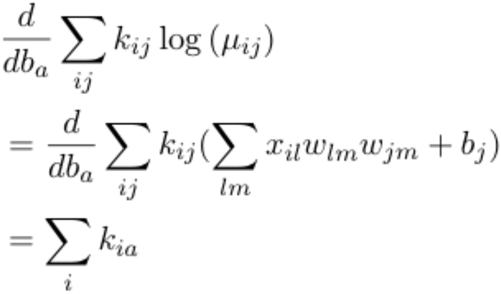

Equivalently for the third therm the derivative is 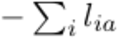 and so the derivative of the loglikclihood with respect to the bias is:

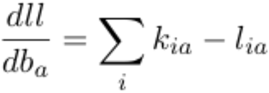

